# Visualising formation of the ribosomal active site in mitochondria

**DOI:** 10.1101/2021.03.19.436169

**Authors:** Viswanathan Chandrasekaran, Nirupa Desai, Nicholas O. Burton, Hanting Yang, Jon Price, Eric A. Miska, V. Ramakrishnan

**Affiliations:** MRC Laboratory of Molecular Biology, Cambridge CB2 0QH, UK; Centre for Trophoblast Research, Department of Physiology, Development and Neuroscience, University of Cambridge, Cambridge, CB2 3EG, UK; Gurdon Institute, University of Cambridge, Cambridge, CB2 1QN, UK; Department of Genetics, University of Cambridge, Downing Street, Cambridge, CB2 3EH, UK; Wellcome Sanger Institute, Wellcome Genome Campus, Cambridge, CB10 1SA, UK

## Abstract

Ribosome assembly is an essential and complex process that is regulated at each step by specific biogenesis factors. Using cryo-electron microscopy, we identify and order major steps in the formation of the highly conserved peptidyl transferase centre (PTC) and tRNA binding sites in the large subunit of the human mitochondrial ribosome (mitoribosome). The conserved GTPase GTPBP7 regulates the folding and incorporation of core 16S ribosomal RNA (rRNA) helices and the ribosomal protein bL36m, and ensures that the PTC base U3039 has been 2′-O-methylated. Additionally, GTPBP7 binds the RNA methyltransferase NSUN4 and MTERF4, which facilitate earlier steps by sequestering H68-71 of the 16S rRNA and allowing biogenesis factors to access the maturing PTC. Consistent with the central role of NSUN4•MTERF4 and GTPBP7 during mitoribosome biogenesis, *in vivo* mutagenesis designed to disrupt binding of their *Caenorhabditis elegans* orthologs to the large subunit potently activates mitochondrial stress responses and results in severely reduced viability, developmental delays and sterility. Next-generation RNA sequencing reveals widespread gene expression changes in these mutant animals that are indicative of mitochondrial stress response activation. We also answer the long-standing question of why NSUN4 but not its enzymatic activity, is indispensable for mitochondrial protein synthesis in metazoans.

## Main

Mitoribosomes are essential for the production and maintenance of the electron transport chain machinery. As with all other ribosomes, the biogenesis of mitoribosomes involves a complex series of coordinated steps during which the ribosomal RNA folds, proteins are incorporated and the active site is formed. The process is facilitated by specific biogenesis factors, including GTPases, rRNA modifying enzymes, RNA helicases, ribonucleases, chaperones and other scaffolding and adaptor proteins (reviewed in^1^). Studying this process in molecular detail is clinically important because defective mitoribosome assembly can lead to encephalomyopathy, optic neuropathy, cardiomyopathy and hereditary spastic paraplegia, among other disorders.

The general principles of ribosome assembly are well-conserved across species, but mitoribosome assembly has diverged in notable ways from and involves different factors than bacterial 70S ribosomes, owing to considerable divergence between bacterial and mitochondrial ribosomes. Analogous to bacterial and eukaryotic cytosolic ribosomes, mitoribosome assembly also proceeds via a series of quality-control checkpoints and GTP hydrolysis serves as the commitment step to drive the reaction forward^2,3^. Four GTPases, GTPBP5,6,7,10 (and perhaps GTPBP8^4^) directly bind the 16S ribosomal RNA (rRNA) of the 39S mitoribosomal large subunit (mtLSU) to facilitate its maturation^5^, but their mechanistic roles and order of action remain poorly understood.

### NSUN4, MTERF4 and MTG1 co-assemble on the subunit interface of the mitoribosomal large subunit

To trap GTPBP7 and other GTPases that participate in human mitochondrial translation, we included the non-hydrolysable GTP analogue β,γ-methyleneguanosine 5′-triphosphate (GMPPCP) during mitoribosome purification (**Methods**). Single particle cryo-EM (Extended Data Fig. 1, 2, Extended Data Table 1) of these and other mitoribosome-bound complexes^6^ also yielded a 5% subset of mtLSU intermediates bound to GTPBP7•GMPPCP in a pre-hydrolysis state and arrested at an assembly state equivalent to the 45S (RI_50_) to 50S transition of the bacterial LSU^7-9^. In this state, the peptidyl transferase centre is not completely disordered as it is in PDB 5OOM^10^, but key helices in the vicinity are held in an unfolded state.

**Fig. 1.**
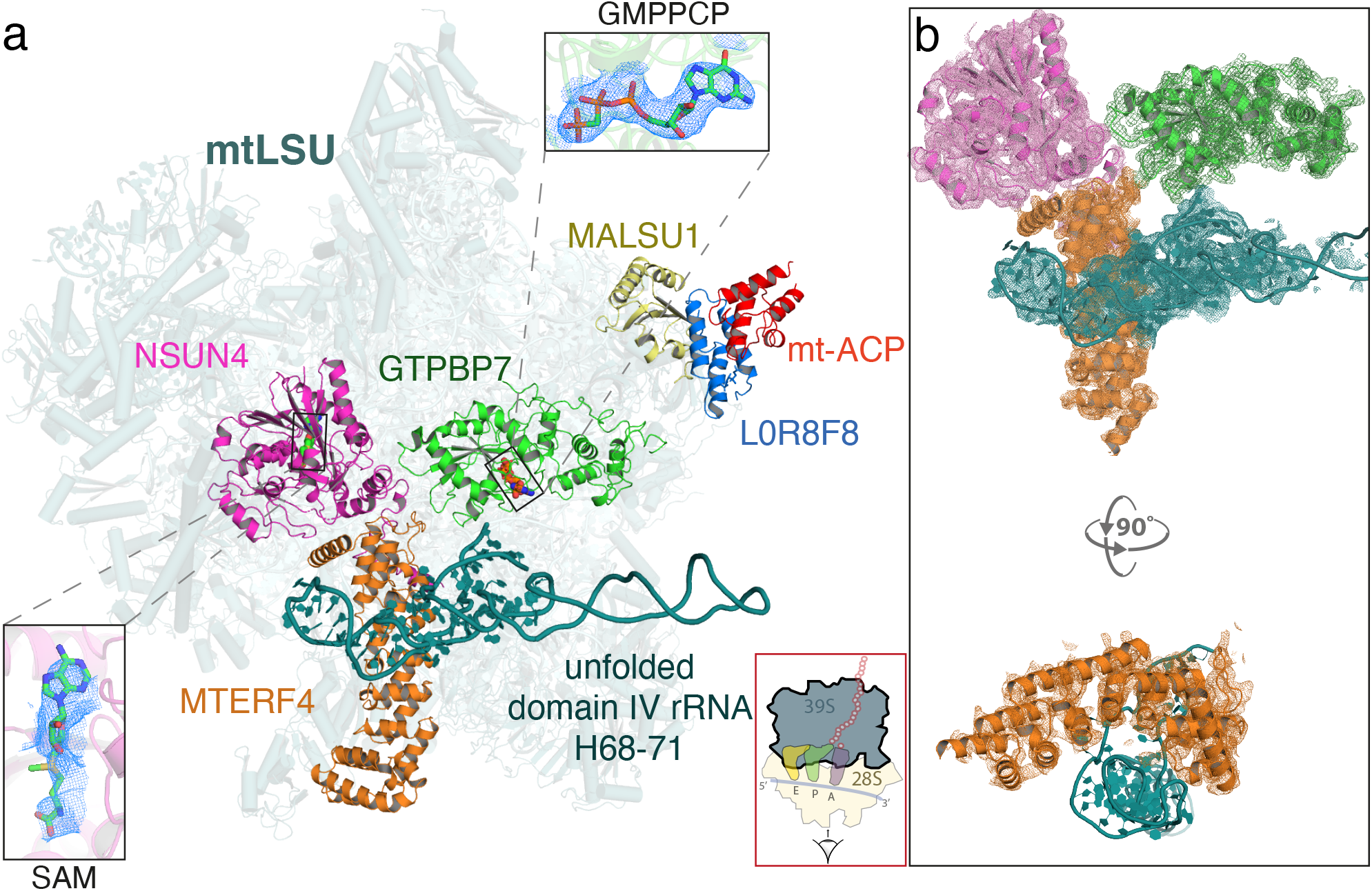
Architecture of the mitoribosomal LSU complexed with NSUN4•MTERF4•GTPBP7•GMPPCP and MALSU1-L0R8F8-mt-ACP. (a) The inter-subunit interface faces the reader and a cartoon of an elongating mitoribosome is shown in the red box (bottom) to aid orientation. NSUN4 (pink) bound to S-adenosyl methionine (SAM, left inset), GTPBP7 (green) bound to the non-hydrolysable GTP analog, GMPPCP (top inset) and MTERF4 (orange) interact with the subunit interface of the mtLSU assembly intermediate. The domain IV rRNA helices H68-71 (teal) are unfolded, rendering the aminoacyl, peptidyl and exit sites incomplete. Also pictured are the anti-association factors MALSU1 (yellow), L0R8F8 (blue) and mt-ACP (red). (b) Map-to-model fits for NSUN4, MTERF4, GTPBP7 and the unfolded rRNA.

Surprisingly, GTPBP7 was bound to a heterodimer of the 5-methyl cytosine (m^5^C) modifying enzyme, NOP2/Sun RNA methyltransferase 4 (NSUN4) and its binding partner, mitochondrial transcription termination factor 4 (MTERF4)^11^ (Fig. 1). NSUN4, MTERF4 and GTPBP7 bind the inter-subunit interface of the mtLSU in a radial spoke arrangement centred about the peptidyl site (P site) and < 30 Å from the peptidyl transferase centre (PTC)^12^. Additionally, the anti-association factors MALSU1•L0R8F8•mt-ACP are also present on these particles to prevent premature SSU joining^6,10^.

The NSUN4 active site contains weak density for the methyl donor S-adenosylmethionine (SAM). Moreover, the cryoEM density suggests that there is no m^5^C methylation of 16S rRNA located within ∼20 Å of the NSUN4 active site indicating that methyltransferase activity is not involved at the mtLSU assembly checkpoint. Bisulphite sequencing studies across many species have also not revealed any m^5^C targets on the 16S rRNA making this an unlikely site of NSUN4 methyl transferase activity^13^.

Our structure represents an intermediate in which the proteins uL16m, bL27m and bL36m have already bound, but before complete folding of rRNA domain IV helices 68-71 and dissociation of NSUN4, MTERF4 and GTPBP7 by GTP hydrolysis, and subsequent mtSSU joining. The 16S rRNA domain IV helices 68-71 (nucleotides 2542-2637) typically span the A, P and E sites and pack against the D-loop, and anticodon arms of the P-site tRNA (Fig. 2). Here, they are instead held in a partially unfolded state by MTERF4 at a location that would permit biogenesis factors access to their binding sites, but clash with an incoming SSU (Fig. 2c). Subsequent GTP hydrolysis by GTPBP7 results in dissociation of the GTPase and MTERF4•NSUN4, allowing H68-71 to fold (compare Fig. 2c, d).

**Fig. 2.**
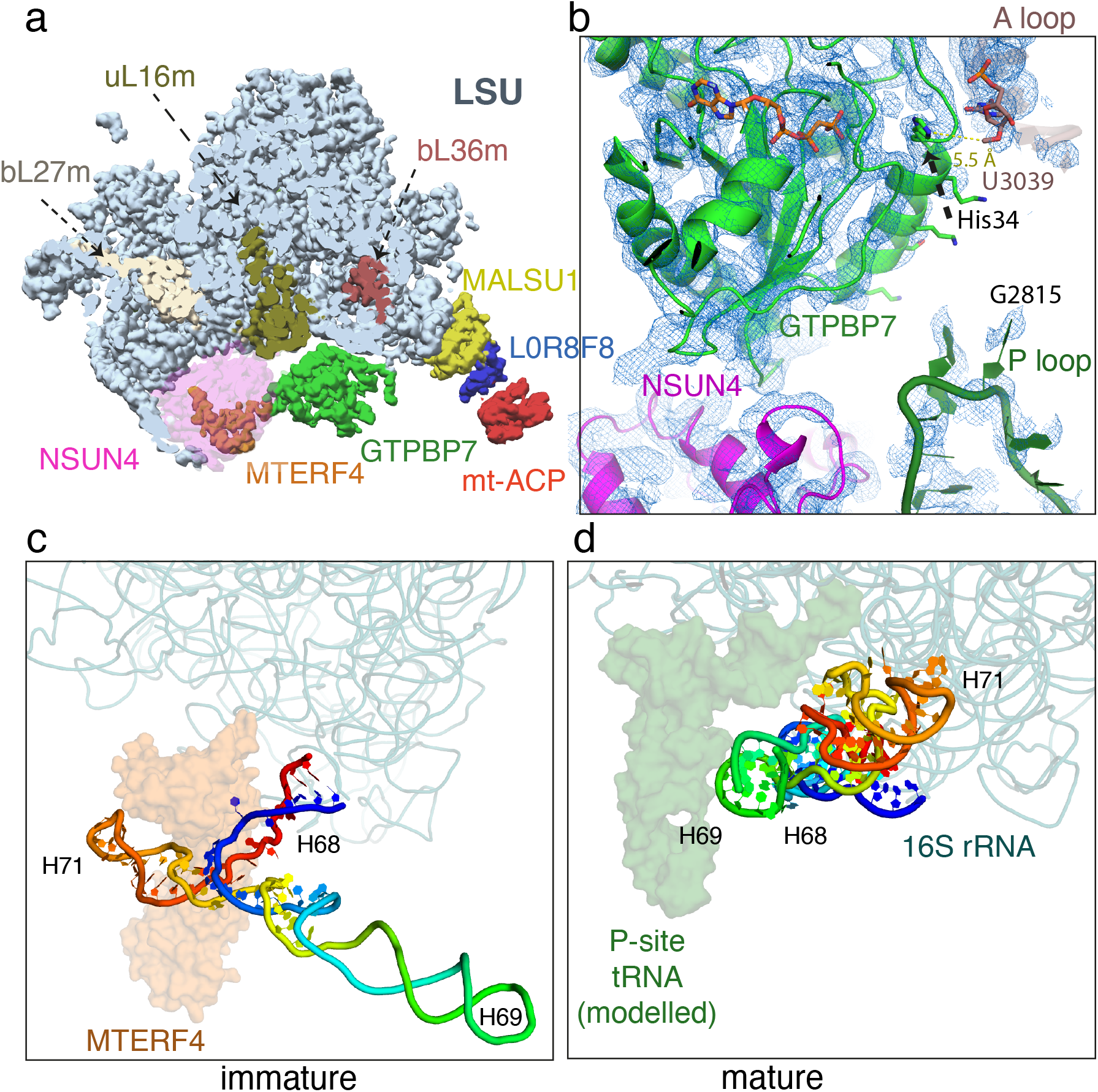
Maturation of the mtLSU by NSUN4•MTERF4•GTPBP7. (a) Incorporation of uL16m (olive), bL27m (wheat) likely precedes NSUN4 (magenta) binding and before bL36m (brown). NSUN4 is shown translucent for clarity. (b) His34 of helix 1 of GTPBP7 is 5.5 from, and ensures 2’-O-methylation of A-loop base U3039 by MRM2. U3040, which is methylated by MRM3 is disordered and not shown in the figure. Immature (c) and mature (d) conformations of domain IV rRNA helices H68-71 (rainbow). H69 would pack against a canonical P-site tRNA when present and together this region forms the back wall of the A, P, and E sites of the mature 39S. MTERF4 (translucent orange) holds H68-71 in an unfolded conformation to permit assembly factors to access the core of the mtLSU.

Analogous to bacterial RbgA, GTPBP7 may facilitate the incorporation of the mitoribosomal protein bL36m into the mtLSU^3,5^ (Extended Data Fig. 3). On the other hand, two other ribosomal proteins also attributed to RbgA, uL16m, and bL27m, are already present in an earlier assembly intermediate (PDB 5OOM). We suspect that this difference is likely because only two GTPases (ObgE being the other) regulate bacterial 50S assembly while at least four are implicated in assembly in humans^4^. Remarkably, GTPBP7 binds in the vicinity of 2’-O-methylated bases in the A-and P-loops of the PTC, which are critical components that bind the aminoacyl and peptidyl tRNAs, respectively. Our structure reveals that the A loop moves towards GTPBP7 (Extended Data Fig. 4) and helix 1 of GTPBP7 directly contacts the highly conserved U3039 to verify 2’-O-methylation by mitochondrial methyltransferase 2 (MRM2)^14^ (Fig. 2b, Extended Data Fig. 4). Given the universal importance of 2’-O-methylation of specific nucleotides in the PTC for translation^15,16^, it is striking that GTPBP7 directly contacts U3039. In principle, slight rearrangements of helix 1 of GTPBP7 could also ensure G3040 methylation by MRM3. Similarly, G2815 (in the P loop), which is 2’-O-methylated by MRM1^14,16^ is in the vicinity of, but is not contacted directly by NSUN4 and GTPBP7 in our structure (Extended Data Fig. 4). G2815 and 3040 are flexible in our map and their methylation status is therefore unknown.

### Destabilizing 39S binding of the three biogenesis factors in *C. elegans* leads to defects in growth and sterility and potently activates mitochondrial stress response

GTPBP7, MTERF4 and NSUN4 are highly conserved and their counterparts in C. elegans are respectively MTG-1, MTERF-4 and NSUN-4. In *C. elegans*, the dispensability of the methyltransferase activity of NSUN4 for mitochondrial protein synthesis was previously characterized^17^, suggesting that its importance may lie in its interaction with the mitoribosomal large subunit. To test the importance of the formation of the complex of the three protein factors with the mtLSU during assembly *in vivo*, we used *C. elegans* as a model organism and introduced mutations at locations in the three factors that we predicted would disrupt binding to the mtLSU (Extended Data Fig. 5).

Mutations that disrupt mitoribosome binding to MTER-4 and MTG-1 (the *C. elegans* orthologs of MTERF4 and GTPBP7 respectively), led to a substantial increase in expression of the mitochondrial heat-shock protein HSP-6, consistent with activation of the mitochondrial unfolded protein response (Fig. 3a). These findings are similar to the effects observed when mitochondrial ribosomal proteins are knocked down by RNAi^18^. By contrast, we found that mutations that disrupt the catalytic activity of NSUN-4 did not substantially activate *hsp-6* expression (Fig. 3a, NSUN-4 cd), nor did mutations that disrupt a weak contact between NSUN-4 and the mitoribosomal protein bL33m. These results are consistent with the catalytic activity of NSUN-4 being dispensable for mitochondrial protein synthesis. However, we note that NSUN-4 makes multiple contacts with the mitoribosome and that the chosen M225A and K226A substitutions in NSUN-4 were likely insufficient to disrupt its interaction with the mitoribosome.

**Fig. 3.**
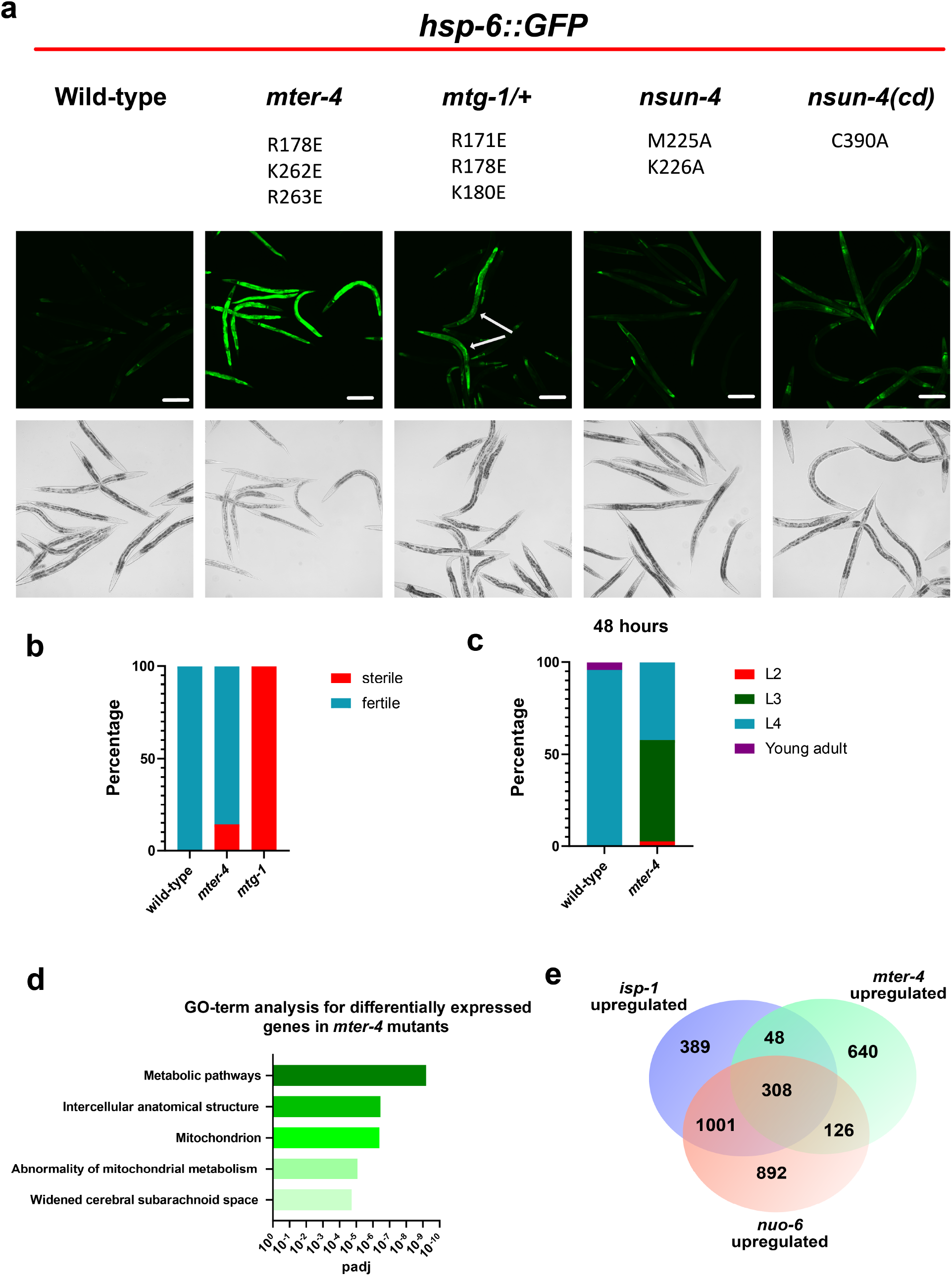
Mutations that disrupt mitoribosomal LSU binding result in activation of the mitochondrial unfolded protein response, delayed development, and decreased fertility. (a) Representative images of *hsp-6::GFP* expression in wild-type, *mter-4(syb3662, syb3403), mtg-1(syb3641)/(s2023) [dpy-1(s2170) umnIs21] III], nsun-4(syb3514)*, and catalytic dead *nsun-4(mj457)* mutant animals. White arrows in mtg-1 mutants represent homozygous mutant animals based on absence of *umnIs21* reporter expression from balancer chromosome. All animals express the *zcIs13 [hsp-6::GFP]* reporter which is specifi-cally expressed in response to mitochondrial stress. Animals were grown for 72 hours at 22.5°C. scale bars = 200 µm (b) Frac-tion of wild-type and *mter-4(syb3662, syb3403)* mutants at different developmental stages after 48 hours at 22.5°C. n = 200 animals (c) Fraction of wild-type and *mter-4(syb3662, syb3403)* mutants sterile at 22.5°C. n = 100 animals (d) Top 5 GO-terms based on p-value from g:Profiler analysis of genes upregulated greater than 0.5 (log2) in *mter-4(syb3662, syb3403)* mutants when compared to wild-type animals. (e) Venn diagram comparison of genes upregulated in *mter-4(syb3662, syb3403), isp-1(qmv150), and nuo-6(qm200)* mutants. *isp-1* and *nuo-6* gene expression data from^20^.

At the organismal level, destabilizing 39S binding by MTG-1 and MTER-4 resulted in delayed animal development and sterility (Fig. 3b, c). Specifically, we found that *mter-4* mutants exhibited delayed development and 14% of mutants were sterile (Figure 3b, c). Similarly, we observed a more severe phenotype in *mtg-1* mutants and found that 100% of homozygous *mtg-1* mutants were sterile (Figure 3b). The severity of the phenotypes of the *mter*-4 and *mtg*-1 mtLSU binding mutants underscores the importance of the role of MTERF4/MTER-4 and GTPBP7/MTG-1 in the mitochondrion. In addition, our results suggest that the effects of disrupting GTPBP7/MTG-1 and MTERF4/MTER-4 interactions with the mitoribosome can be overcome in somatic tissues, potentially by activating the mitochondrial unfolded protein response, as mutant animals can indeed develop to adulthood. Germ cells, however, were particularly sensitive to these mutations, and disruption of mtLSU binding caused sterility. Notably, mutations in mitochondrial translational components, including mitochondrial tRNAs, are known to cause sterility in humans (reviewed in ^19^). We suggest that germ cell sensitivity to disruptions in mitochondrial translation is a conserved phenomenon throughout metazoans and establish a new model to study these effects.

To further confirm the effects of disrupting the interaction of these proteins with the mitoribosome, we performed next-generation RNA sequencing on the viable *mter-4* mutants. We identified 3,174 differentially expressed genes in *mter-4* mutants when compared to wild-type animals (Extended Data Table 2). Furthermore, we found that the genes upregulated in *mter-4* mutants are enriched for genes associated with mitochondrial dysfunction and substantially overlap with genes that are upregulated in electron transport chain mutants^20^ (Fig. 3d, e). These results support our structural data and further suggest that disrupting the interaction of these proteins with the mitoribosome activates the mitochondrial unfolded protein response and disrupts normal mitochondrial function. The pathological significance of these proteins is demonstrated by their association with cardiomyopathy, which is a common feature of mitochondrial diseases. Cardiomyocyte-specific MTERF4 knock-out (KO) mouse models develop mitochondrial cardiomyopathy, and global KO mice are embryonically lethal^11^. Variant alleles of MTERF4 have also been documented in a paediatric patient with hypertrophic cardiomyopathy^21^. Similarly, GTPBP7 and NSUN4 silencing in human cardiomyocytes^22^ and conditional KO mice have been shown to lead to impaired cardiac physiology and progressive cardiomyopathy^13^, respectively.

Our observations of severe mitochondrial defects and activation of the mitochondrial unfolded protein response in *mter-4* mutants, but not *nsun-4* catalytic dead mutants, suggests that the major function of these proteins is as a checkpoint in mitochondrial ribosome assembly and not in RNA methylation. Consistent with this idea, MTERF4 does not participate during NSUN4-mediated methylation of the decoding centre of the mtSSU^11^. We have thus shown, using structure-directed mutagenesis *in vivo*, that while the methyltransferase activity of NSUN4 on the small subunit can be dispensed with without resulting in a strong phenotype, the protein has a more important non-enzymatic function as a biogenesis factor in the final stages of the formation of the PTC and the tRNA binding sites in the large subunit.

### A Scheme for the assembly of the peptidyl transferase centre

We have previously reported the cryo-EM structures of two human late-stage mitoribosomal assembly intermediates^23^. These structures when taken together permit piecing together the order of events during late-stage maturation of the mtLSU. The anti-association factors MALSU1, L0R8F8 and mt-ACP engage the mtLSU at an earlier stage of assembly since they are present even when the entire domain IV (H68-71) is disordered^23^ (Fig. 4, state *I*). At this stage, ribosomal proteins bL27m and uL16m have already been incorporated and at some point, are followed by bL36m. In a later stage of assembly, the binding of MTERF4 binds to and holds open H68-71 in an unfolded state, while NSUN4 binds to the inter-subunit interface. GTPBP7•GTP, whose binding depends on the prior 2′-O-methylation of G2515, U3039 and G3040 that form parts of the A and P loops of the peptidyl transferase centre, binds directly to H34-35, H65-67 and helps fold H89-93. The protein also makes interactions with NSUN4 and MTERF4. Together, all three proteins facilitate correct assembly of the PTC and tRNA sites. GTPBP7 may dissociate following GTP hydrolysis, eject NSUN4-MTERF4 and permit H68-71 to fold into the PTC. MALSU1•L0R8F8•mt-ACP finally leaves the mature 39S that is now ready for subunit joining during initiation. The relative order of dissociation of GTPBP7 and the anti-association factors cannot yet be ascertained, and may actually occur in concert^4,22^.

**Fig. 4.**
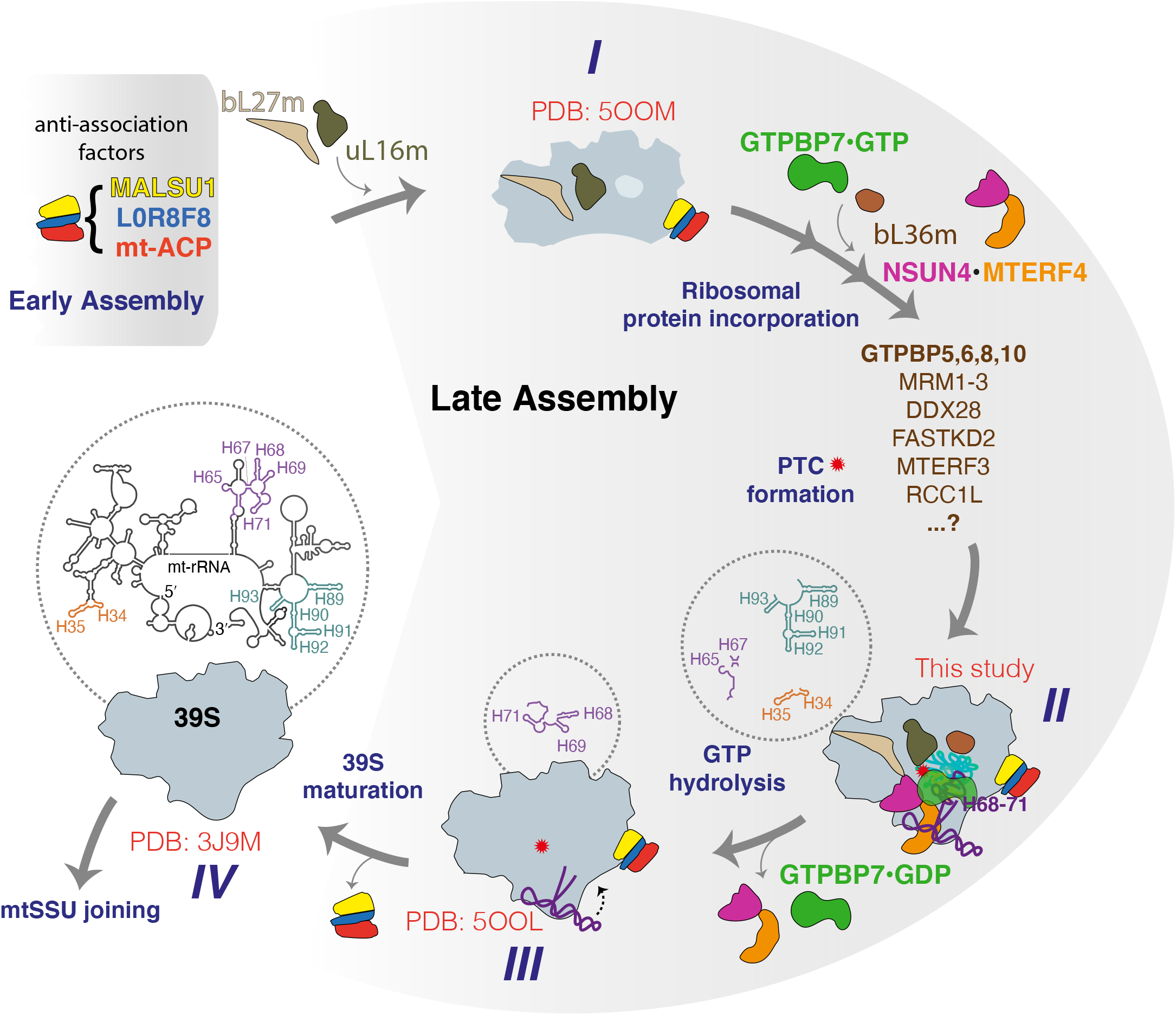
Late events during mitoribosomal large subunit assembly. Schematic of events and their postulated order. The anti-association factors MALSU1, L0R8F8 and mt-ACP as well as the ribosomal proteins bL27m and uL16m, whose incorporation has been attributed to RbgA in bacteria are already present in state I. bL36m is loaded and methylases that target PTC bases and other biogenesis factors that assist rRNA folding act between states I and II. GTPBP7 engages the PTC to verify the methylation status of the A and P loop bases in the PTC. All rRNA elements (except H68-71) are fully folded in II. The biogenesis factors, includ-ing MTERF4 dissociate (likely upon GTP hydrolysis by GTPBP7) to allow H68-71 to also fold to reach state III. MALSU1, L0R8F8 and mt-ACP dissociate and the mature 39S is formed IV.

The proper folding of the peptidyl transferase centre, the universally conserved active site of the ribosome, along with the tRNA binding sites, is one of the crucial steps in the assembly of the large subunit. We have shown that this assembly occurs in multiple steps, moving from one in which the elements of the PTC are in a highly disordered state to one where they are ordered but not yet finally folded into their final structure in the large subunit. This assembly requires several protein factors to act together to yield the mature ribosomal subunit including factors previously unidentified as assembly factors. By coupling cryo-EM analysis of human mitoribosome assembly intermediates with *in vivo* studies in *C. elegans* we have identified the physiological importance of these factors in an essential, conserved late-stage mtLSU maturation checkpoint. Although our studies were specifically on mitoribosomes, given the universally conserved nature of the PTC and its importance to the ribosome, we expect that other domains of life will have a similarly complex and orchestrated process requiring multiple factors to facilitate the formation of the active sites of the large ribosomal subunit.

## Methods

### Cell Line

PDE12^-/-^ HEK293T cells that were used have previously been published^24^. Initially cell growth was adherently at 37°C, 5% CO_2_, in 10% fetal bovine serum (FBS) supplemented Dulbecco’s Modified Eagle Medium (DMEM). Subsequently the cells were adaptation to be grown in suspension in 1% FBS supplemented Freestyle media at 37°C, 8% CO_2_.

### Purification of native mitoribosomal complexes

Human mitochondria, and from them native mitoribosomal complexes, were purified from PDE12^-/-^ HEK293T cells as previously described^6^. Briefly, purified cells were harvested at 10^6^ cells/ml and washed in cold phosphate buffered saline (PBS). The weighed pellet of cells was resuspended in 6 ml of MIB buffer (50 mM HEPES-KOH pH 7.5, 10 mM KCl, 1.5 mM MgCl_2_, 1 mM EDTA, 1 mM EGTA, 1 mM DTT, Proteinase Inhibitor; PI, 1 tablet per 50 ml) per gram and SM4 buffer (281 mM sucrose, 844 mM mannitol in MIB buffer) to yield 70 mM sucrose and 210 mM mannitol. The cells were subjected to nitrogen cavitation for 20 minutes at 500 psi to disrupt the cell membranes and the collected sample was centrifugation twice at 800 x g for 15 minutes and the supernatant collected. Two additional centrifugation steps at 10,000 x g for 15 minutes were performed and each time, the pellet was resuspended in 0.5mls of MIBSM buffer (3:1 MIB:SM4) per 1g of originally weighed pellet. Per 1g of the original pellet 10U of RNase free DNase was added and rotated on a roller at 4°C for 20 minutes. The solution was then pelleted at 10,000 x g for 15 minutes, resuspended in SEM buffer (250 mM sucrose, 20 mM HEPES · KOH pH 7.45, 1 mM EDTA), dounce-homogenised, and layered onto a sucrose step gradient of 60%, 32%, 23%, 15% sucrose for centrifugation at 99,004 x g for 1 hour at 4°C. The collected mitochondrial layer was flash frozen in liquid nitrogen for storage at -80°C.

To isolate mitoribosomes from the collected mitochondrial fraction, 2 volumes of lysis buffer (25 mM HEPES.KOH pH 7.4, 100 mM KCl, 25 mM Mg(OAc)_2_, 1.5% β-DDM, 0.15 mg/ml TOCL, 0.5 mM GMPPCP (Sigma Aldrich), 1 tablet per 50 ml PI, 2 mM DTT) were added and briefly dounce-homogenized and stirred for 30 minutes in the cold room. Supernatant was collected after centrifugation at 30,000 x g for 30 minutes and loaded at a ratio of 2.5:1 onto a 1M sucrose cushion (20 mM HEPES.KOH, pH 7.4, 100 mM KCl, 20 mM Mg(OAc)_2_, 0.6% β-DDM, 0.06 mg/ml TOCL, 0.25 mM GMPPCP, 2 mM DTT) and centrifuged at 231,550 x g for 60 minutes at 4 °C. The pellet was resuspended (20mM HEPES.KOH, pH 7.4, 100 mM KCl, 5 mM Mg(OAc)_2_, 0.3% β-DDM, 0.03 mg/ml TOCL, 0.25 mM GMPPCP, 2 mM DTT) and the crude mitoribosome fraction was layered onto a 15-30% sucrose gradient and subjected to centrifugation at 4°C for 90 minutes at 213,626 x g. Following sucrose gradient fractionation, the mitoribosomal fractions were pooled and concentrated into final buffer (20 mM HEPES.KOH, pH 7.4, 100 mM KCl, 5 mM Mg(OAc)_2_, 0.05% β-DDM, 0.005 mg/ml TOCL, 0.25 mM GMPPCP, 2 mM DTT) before the mitoribosomes were vitrified for cryo-EM.

### Grid preparation and Data collection

Quantifoil R2/2 holey carbon grids, covered with home-made amorphous carbon (∼50 Å thick) was used with prior glow discharging. Using the Vitrobot Mk IV, 3 μl of our sample was applied to the grids, blotted for 4-6 seconds, vitrified by plunging into liquid ethane and stored in liquid nitrogen. Data was collected over 7 separate sessions in in integrating mode at a magnification of 75,000, pixel size 1.04 Å on the FEI Titan Krios 300 kV electron microscope using the FEI Falcon III detector and EPU software. A total of 46,109 movies were collected using a defocus range from -1.1 to -3.2 at a dose of 1.5 e^−^ per frame per Å^2^ at 1 s exposure (39 frames).

### Image Processing

RELION-3.0 and 3.1^25^ was used to process cryo-EM data as described before^6^. Resolutions are reported according to the Fourier shell correlation (FSC) = 0.143 criterion^26^. Movie motion correction was performed using MotionCorr2 and the contrast transfer function was estimated using CTFFIND-4.1^27^. 3,374,367 particles were picked from 43,950 micrographs using a 2D generated reference (Extended Data Fig. 1). Micrographs with CTF figure of merit >0.3 and a maximum resolution better than 5 Å were selected for further processing. Following particle extraction (128-pixel box; 5.0375 Å/pixel) and 2D classification, 3,247,481 particles were retained. The first 3D refinement yielded a 10.2 Å map at Nyquist resolution, using a 60 Å lowpass-filtered reference mitoribosome. The reported 3D refinements are based on gold-standard estimates. A general 3D classification without alignments separated out mtLSU subclasses consisting of 1,283,454 particles. Following refinement and further 3D classification without alignment, a class of 1,100,599 particles was obtained that had previously unmodeled cryo-EM density at the subunit interface. This class was re-extracted to a pixel size of 1.04 Å and refined to 3.1 Å. Two subsequent rounds of focussed classification with signal subtraction (FCwSS) were performed masking the P and E sites and L1 stalk and A, P and E sites respectively. Following 3D refinement, CTF refinement and Bayesian polishing, the LSU assembly intermediate class was refined to 3.4 Å.

### Model building, refinement and validation

PDB 7A5F^6^ was used as a starting model for model building in Coot v 0.9.3^28^. Real-space refinement and validation was performed using phenix.real_space_refine^29^ and the Phenix suite^30^, respectively. Homology models for MTERF4, NSUN4 and GTPBP7 were generated using trRosetta^31^. The unfolded rRNA helices 68-71 were modelled using RNAcomposer^32^ using secondary structure predictions from RNAfold^33^. Only the backbone coordinates but not the bases were retained for the unstructured regions of H68-71. Comparisons were made with PDB 5OOL and 5OOM^23^.

### *C. elegans* alleles and strain maintenance

All strains were grown and maintained at 20 °C on NGM agar plates seeded with *E. coli* HB101 unless otherwise stated. Individual point mutations in *mter-4(syb3662, syb3403), mtg-1(syb3641)*, and *nsun-4(syb3514)* were generated by SunyBiotech (Fuzhou, China). *nsun-4(syb3514)* results in an M225A and K226A substitution in NSUN-4. *mtg-1(syb3641)* results in an R171E, R178E, and K180E substitution in MTG-1. *mter-4(syb3403)* results in an R178E substitution in MTER-4. *mter-4(syb3662)* results in an K262E and R263E substitution in MTER-4. *nsun-4(mj457)* results in a C390A catalytic dead conversion in NSUN-4 and was reported previously^34^. All mutants were crossed with SJ4100 -*zcIs13[hsp-6::GFP] -*to generate *hsp-6* reporter strains. CGC32 -*sC1(s2023) [dpy-1(s2170) umnIs21] III] -*was used to balance *mtg-1(syb3641)* which resulted in sterility.

### *C. elegans* Imaging

Animals were collected as embryos and grown on NGM agar plates seeded with *E. coli* HB101 at 22.5 °C. Adult animals were collected and immobilized in tetramisole and imaged using a Leica DM6 B and a Leica DFC9000 GT camera.

### RNA-seq

Animals were collected as embryos and grown on NGM agar plates seeded with *E. coli* HB101 at 20 °C. Young adults were collected and washed three times in M9 buffer and snap frozen in liquid nitrogen. Pellets of animal tissue were frozen and thawed five times and then refrozen at -70 °C. RNA extraction and sequencing were performed by BGI Genomics (Hong Kong, China). Briefly, RNA was extracted by phenol-chloroform extraction. mRNA molecules were purified from total RNA using oligo(dT)-attached magnetic beads. cDNA was generated using random hexamer-primed reverse transcription. Libraries for paired end 100 bp DNBSeq were generated and validated on the Agilent Technologies 2100 bioanalyzer.

### RNA-seq data analysis

Raw reads were trimmed for adaptors, low quality sequences and short reads with Trimmomatic^35^ (version 0.39, parameters: ILLUMINACLIP:TruSeq3-SE.fa:2:30:10 SLIDINGWINDOW:4:20 MINLEN:20). From the trimmed reads, the remaining ribosomal RNA was removed with sortmeRNA^36^ (version 2.1, default parameters). Clean reads were then mapped to the *C. elegans* reference genome (WBCEL235) with HISAT2^37^ (version 2.1.0, default parameters) and raw counts for each were produced with HTSeq-count^38^. Other quality control metrics were obtained with fastQC, Picard Tools and multiQC^39^. Counts were imported into R and differential gene expression analysis was performed with DESeq2 (FDR < 0.01, LFC > | 0.5 |)^40^. Gene ontology analysis of DEGs was performed with gProfiler^41^.

### Developmental rate and sterility assays

Animals were collected as embryos and grown on NGM agar plates seeded with *E. coli* HB101 at 22.5 °C. To assay developmental rate 200 animals were scored for their developmental stage at 48 hours. To assay for sterility 100 L4 stage animals were each individually transferred to a new plate. Animals with progeny after four days were scored as fertile. Animals with no progeny after four days were scored as sterile.

### Figure Preparation

UCSF Chimera^42^, Pymol (Schrödinger, LLC) and Geneious 10.2.6 (https://www.geneious.com) were used for figure preparations.

## Data availability

Cryo-EM movies will be deposited to the EMPIAR database.

Coordinates and maps will be deposited to the PDB and EMDB respectively.

RNA-seq data is available through NCBI GEO using the accession code GSE169089.

## Acknowledgements

We thank J. Grimmett and T. Darling for advice, data storage, and high-performance computing; M. Minczuk for providing the experimental cell line; the Ramakrishnan lab members for useful discussions and reagents and A. Yerra for discussions and writing support. We acknowledge the MRC Laboratory of Molecular Biology Electron Microscopy Facility for access and support of electron microscopy, sample preparation, and data collection. This work was supported by the UK Medical Research Council MC_U105184332, a Wellcome Trust Senior Investigator award (WT096570), the Agouron Institute, and the Louis-Jeantet Foundation to V.R., and Cancer Research UK (C13474/A18583, C6946/A14492) and the Wellcome Trust (219475/Z/19/Z, 092096/Z/10/Z) to E.A.M. N.D. is funded by a Wellcome Trust Clinical PhD Fellowship (110301/Z/15/Z). NB is funded by a Next Generation Research Fellowship at the Centre for Trophoblast Research. H.Y. is funded by an EMBO Long-term Fellowship (EMBO ALTF 806-2018). For the purpose of Open Access, the author has applied a CC BY public copyright licence to any Author Accepted Manuscript version arising from this submission.

---

**Extended Data Table 1.**
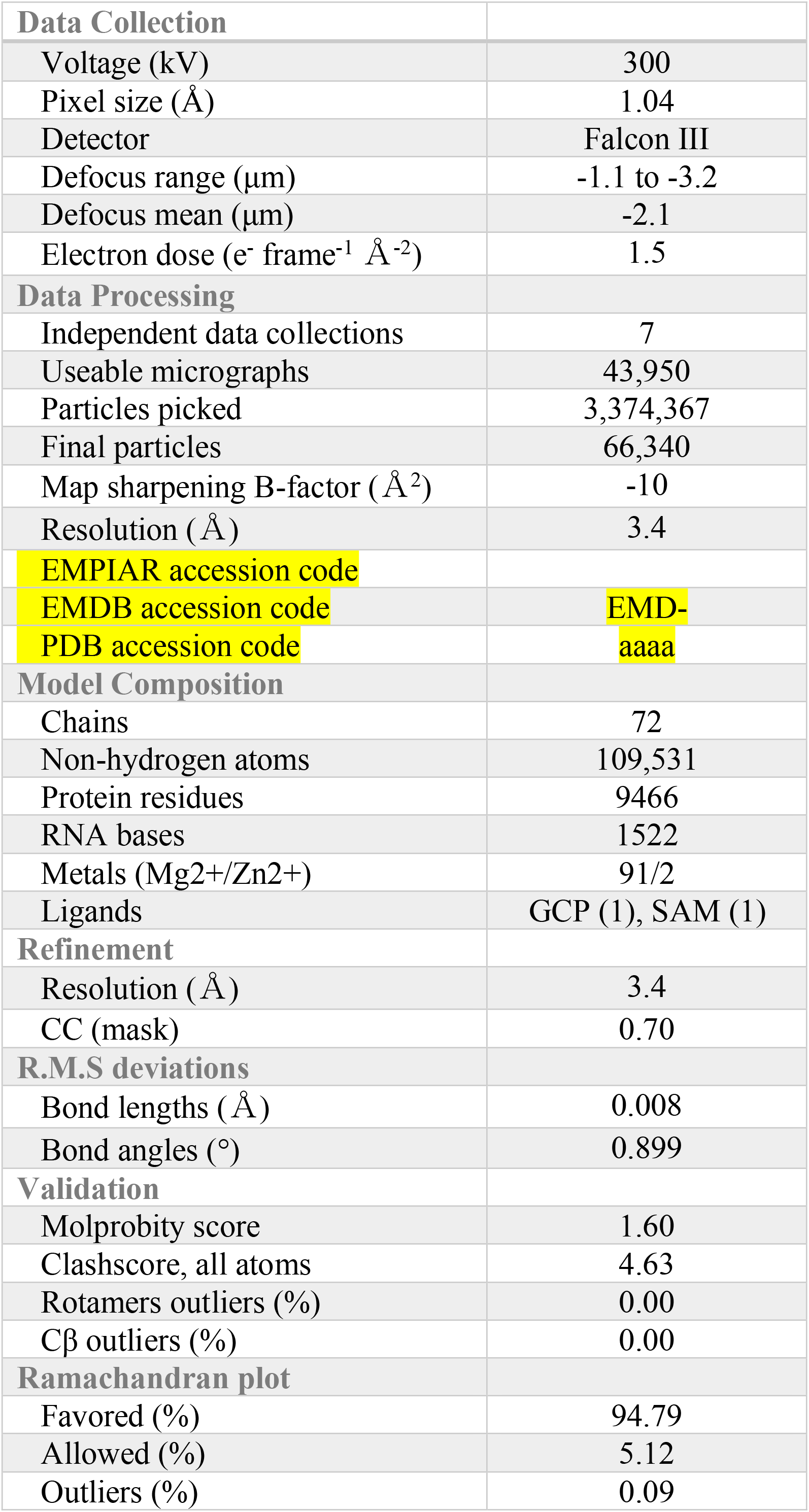
Data collection, processing, refinement and model statistics.

**Extended Data Fig. 1.**
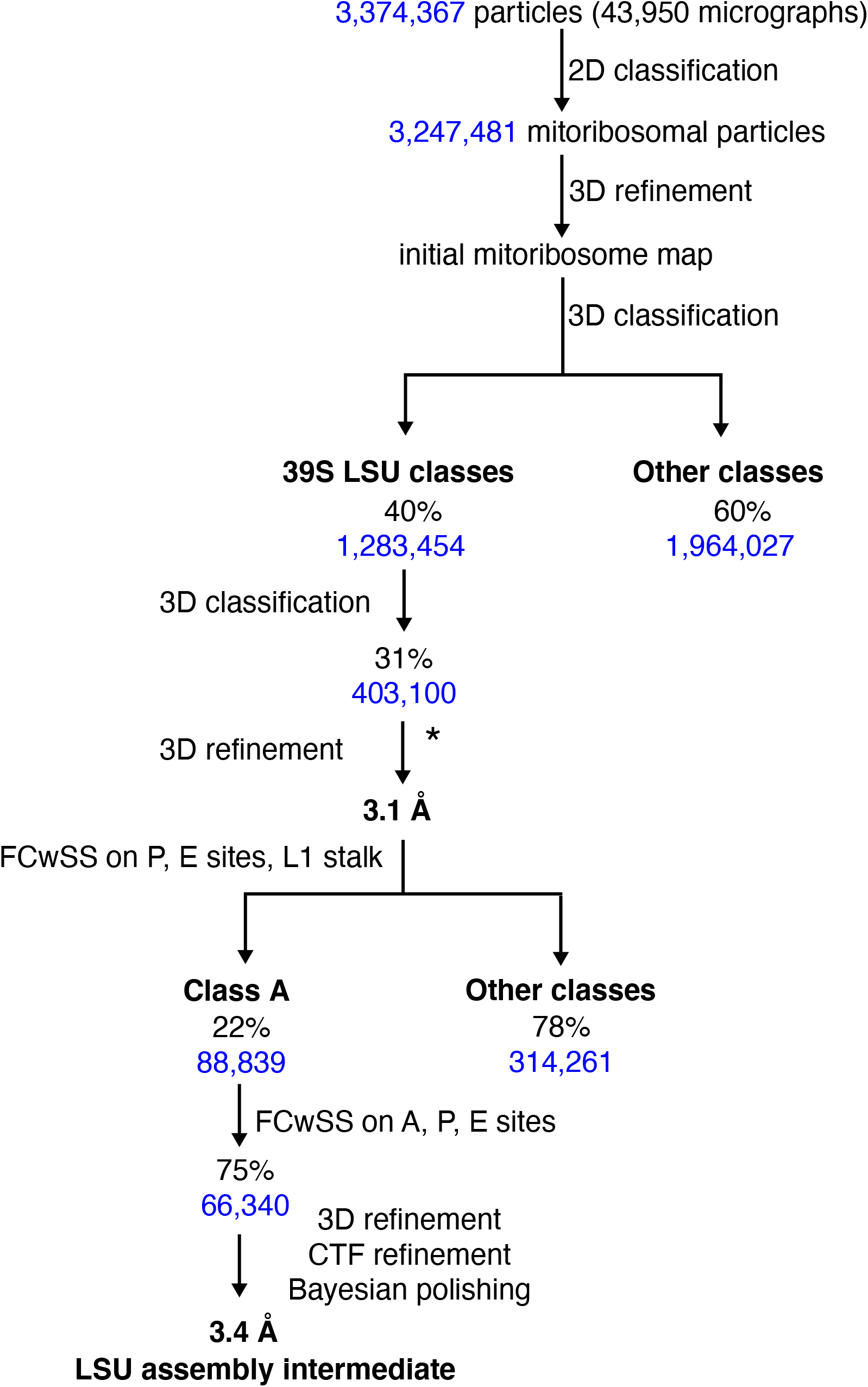
Cryo-EM analysis of human mitoribosomes. Data processing scheme. * indicates that the particles were re-extracted to 1.04 Å/pixel.

**Extended Data Fig. 2.**
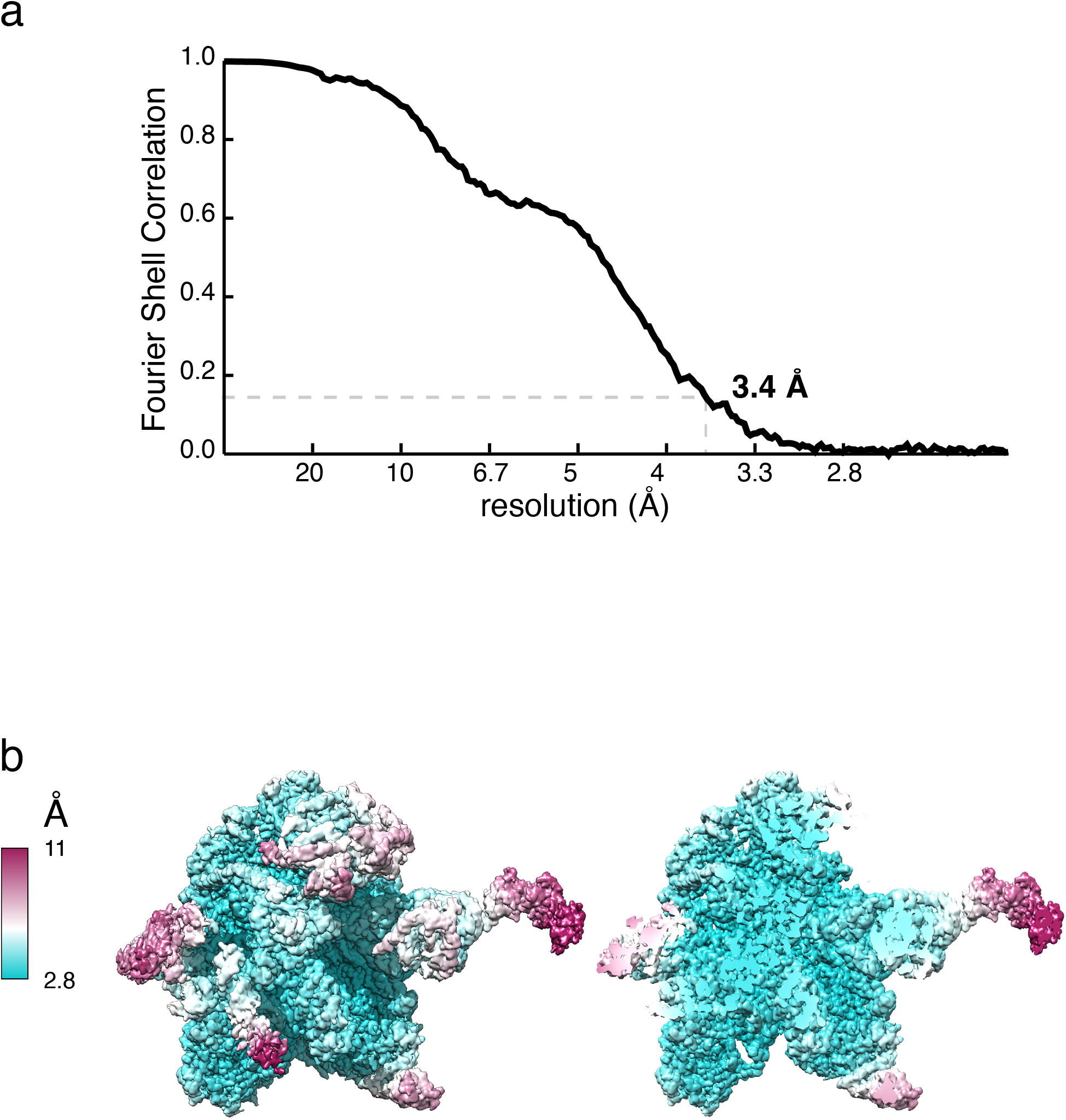
Gold-standard FSC curve (a) and local resolution (b) of the mtLSU assembly intermediate map.

**Extended Data Fig. 3.**
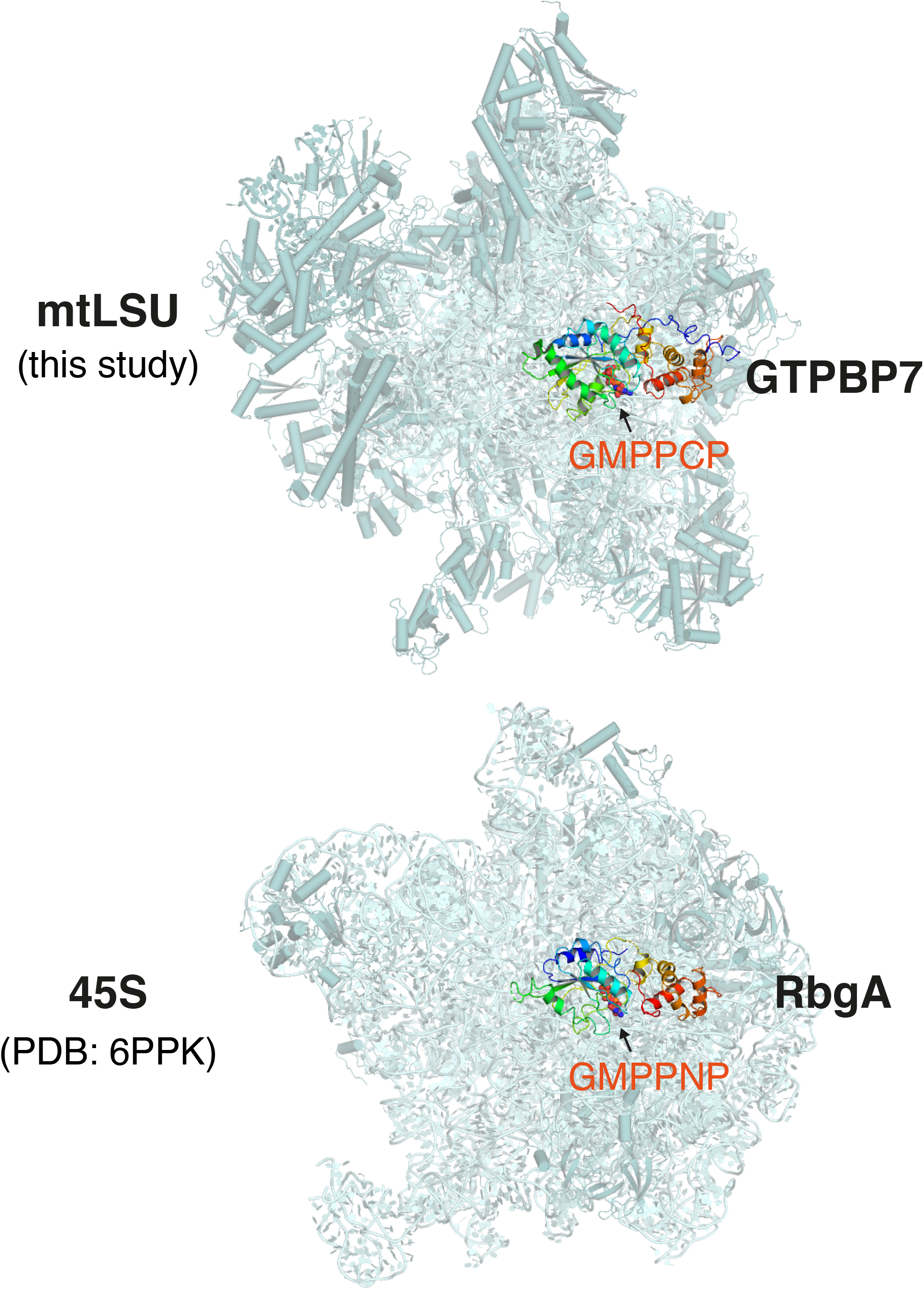
Comparison of MTG1-bound human mtLSU and RbgA-bound *B*.*subtilis* 45S. The inter-subunit interface is oriented towards the reader. GTPBP7 (top) or RbgA (bottom) bound to the non-hydrolysable GTP analogs, GMPPCP (top) or GMPPNP (bottom) interact at equivalent locations on the human mtLSU and bacterial 45S, respectively.

**Extended Data Fig. 4.**
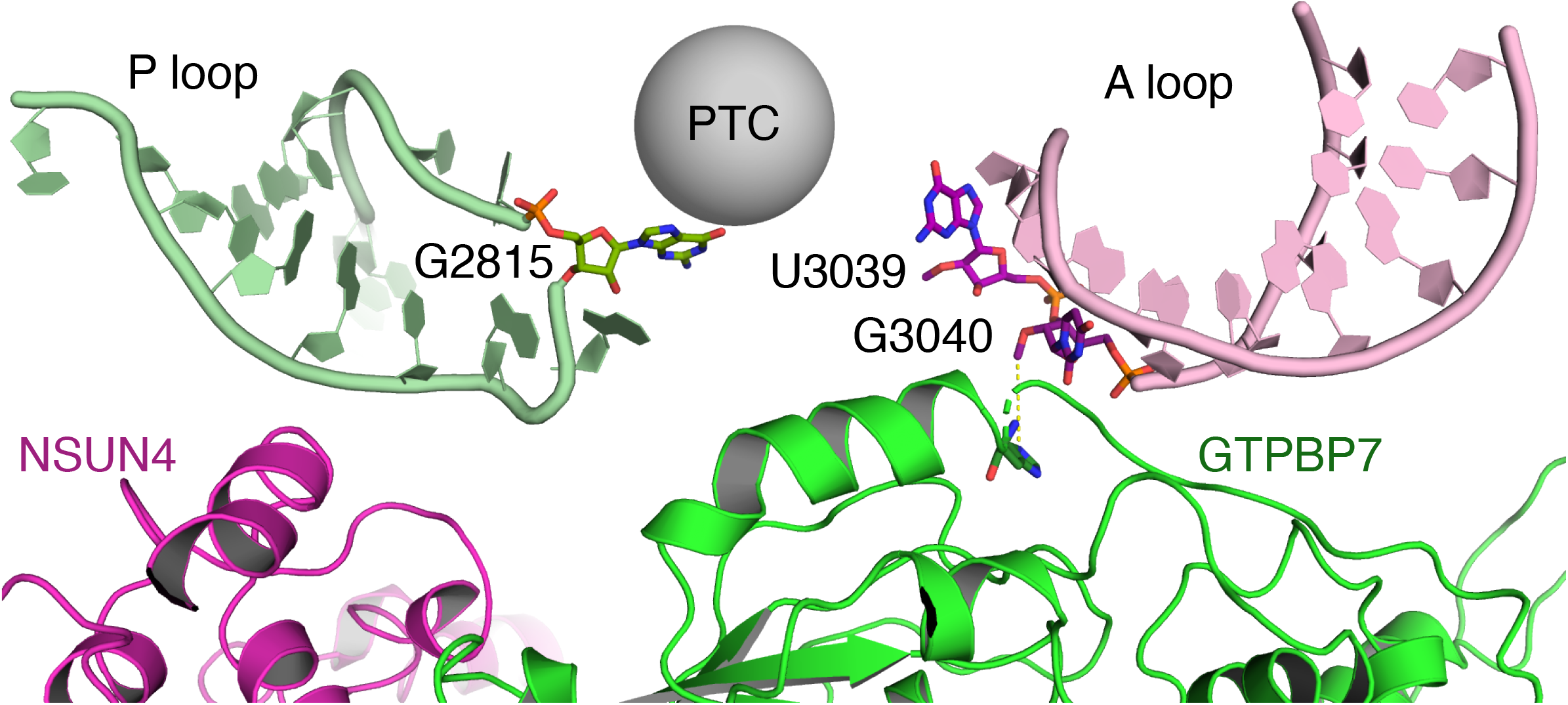
GTPBP7 binds directly to the A loop of the 16S rRNA

**Extended Data Fig. 5.**
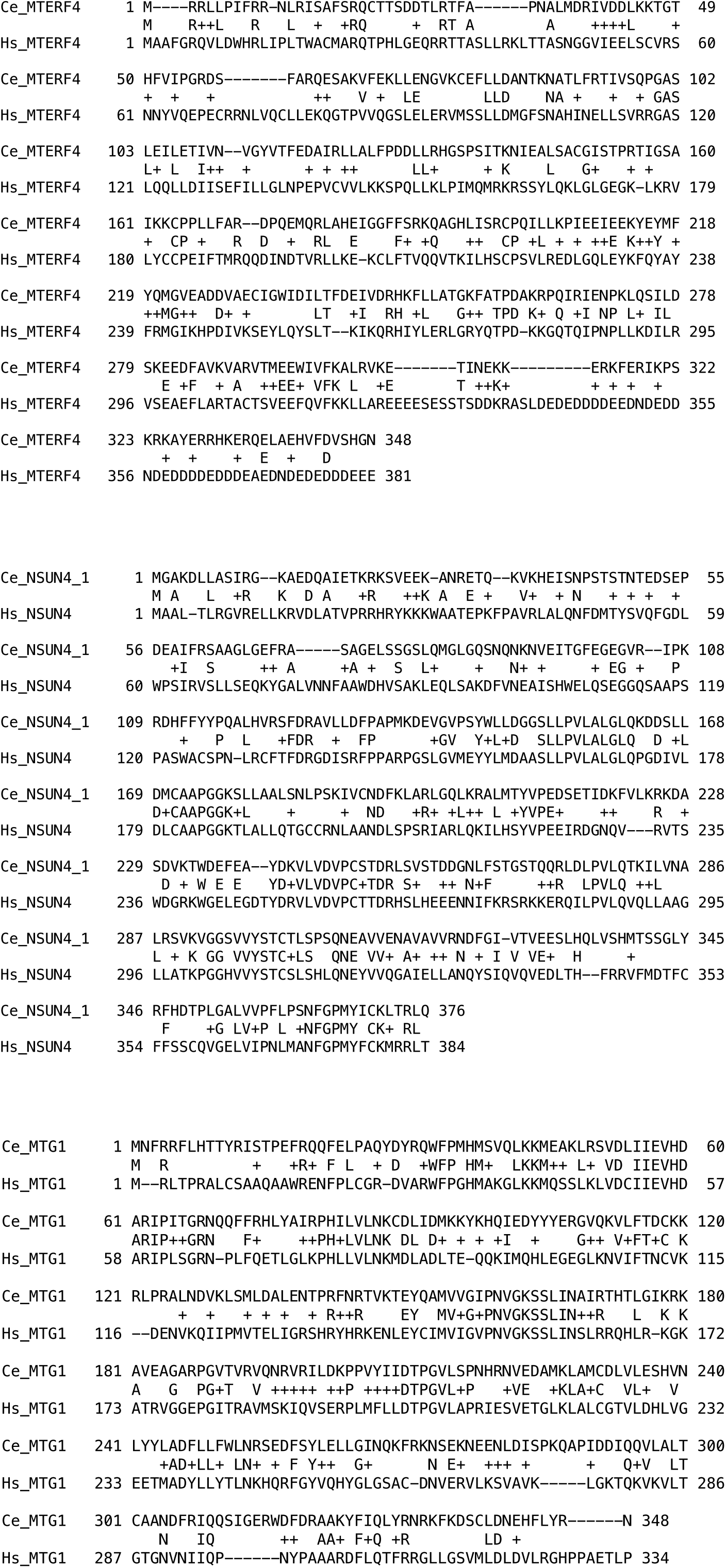
Sequence alignments of C. elegans (Ce) and human (Hs) MTERF4 (top), NSUN4 (middle) and GTPBP7 (MTG1, bottom)

